# A rapid ELISA platform with no sample preparation requirement

**DOI:** 10.1101/2023.04.14.536923

**Authors:** Nicolò Maganzini, Agnes Reschke, Alyssa Cartwright, Yasser Gidi, Ian A. P. Thompson, Amani Hariri, Constantin Dory, Yael Rosenberg-Hasson, Jing Pan, Michael Eisenstein, Timothy Thomas Cornell, Hyongsok Tom Soh

## Abstract

Since its invention in the 1970’s, the enzyme-linked immunosorbent assay (ELISA) has served as the “gold-standard” for blood and plasma protein biomarker quantification. However, ELISAs require significant amounts of sample preparation entailing multiple reagent additions, incubations, and washing steps, limiting their clinical usefulness in the context of diagnosis and prognosis of rapidly evolving medical conditions. In this work, we describe the ‘instant ELISA’ biosensor platform, a probe that can be exposed directly to blood or other biological samples and quantifies protein biomarkers within 15 minutes. The sensor leverages a novel affinity reagent termed ‘monolithic dual-antibody clamp’ (MDAC) which preserves the specificity, sensitivity, and generalizability of ELISA while also enabling rapid analysis of unprocessed blood and other complex matrices. Using MDAC in chicken media, we demonstrate picomolar quantification of the inflammatory marker tumor necrosis factor alpha (TNFα), as well as monocyte chemotactic protein (MCP)-1, a useful prognostic indicator of cytokine release syndrome (CRS) during chimeric antigen receptor (CAR) T-cell immunotherapy. Finally, we demonstrate MCP-1 quantification in plasma samples from patients who had undergone CAR T-cell treatment.

## INTRODUCTION

Since its invention in the 1970’s, the enzyme-linked immunosorbent assay (ELISA) has served as the “gold-standard” for the detection and quantitation of protein biomarkers in blood or other clinical samples.^1^ For example, ELISAs are commonly employed in the context of cancer^2^ (*e.g.*, prostate specific antigen, carcinoembryonic antigen), neurological disease^3^ (*e.g.*, beta-amyloid), and cardiovascular disorders^4^ (*i.e.*, cardiac troponin, creatine kinase). However, ELISAs require significant amounts of sample preparation entailing multiple reagent additions, incubations, and washing steps.^5^ As such, diagnostic ELISAs are typically performed in centralized analytical laboratories by technicians, introducing delays that can potentially impact patient outcomes. This latter concern is particularly relevant in the context acute clinical care for rapidly-evolving medical conditions where timely response is critical.^6–15^

Several groups have attempted to address this need with miniaturized immunoassay platforms that can potentially be applied at the point of care, including systems based on photonic,^16–18^ electrochemical,^19,20^ or field-effect^21,22^ technologies. For example, Song *et al.*^18^, have demonstrated a digital ELISA platform—similar to the commercial SIMOA assay by Quanterix^23^—that they integrated into a microfluidic system and which enables rapid sample-to-answer measurements. Hu *et al.*^19^, adapted a conventional proximity immunoassay to generate an electrochemical readout, a platform that is highly amenable to miniaturization. However, the solutions described to date either cannot operate directly in whole blood or make use of complex microfluidic systems that involve on-chip reagent addition and wash steps. Other approaches have leveraged label-free plasmonic methods^24–26^ to simplify the assay, but this often comes at a cost to the sensitivity, specificity, or generalizability of the assay. Thus, there remains a need for a broadly applicable platform that retains the robust analytical performance of the gold-standard ELISA format while also enabling rapid and direct analysis of patient samples at the point of care.

In this work, we describe the ‘instant ELISA’ biosensor platform, which preserves the specificity, sensitivity, and generalizability of ELISA while also enabling rapid, direct analysis of unprocessed blood. Our sensor makes use of a ‘monolithic dual-antibody clamp’ (MDAC) construct, a novel affinity reagent design that is in turn coupled to the surface of a tapered fiber-optic probe. This probe can be exposed directly to blood or other biological samples and can detect the binding state of the MDAC within 15 minutes. As an initial demonstration, we generated an MDAC for tumor necrosis factor alpha (TNFα), and demonstrated that we can achieve sub-nanomolar sensitivity for the detection of this analyte, even in the complex medium of chicken serum. We subsequently demonstrate the generalizability of this design by generating an MDAC for the detection of monocyte chemotactic protein (MCP)-1, which we subsequently incorporated into the fiber-probe-based instant ELISA assay format, achieving a limit of detection (LOD) of 126 pM in whole chicken blood and 56 pM in chicken plasma. MCP-1 protein is a useful prognostic indicator of cytokine release syndrome (CRS), a life-threatening inflammatory complication that can arise during chimeric antigen receptor (CAR) T-cell immunotherapy. Using the instant ELISA platform, we demonstrate quantitative detection of MCP-1 in the plasma samples from patients who had undergone CAR T-cell treatment within 15 minutes – with no sample preparation.

## RESULTS AND DISCUSSION

### Design of the instant ELISA immunoassay

Instant ELISA (**Figure 1a-d**) integrates multiple technological advances to eliminate the need for the sample preparation and wash steps associated with conventional immunoassays, offering a simpler and faster alternative for analyte detection in complex biological samples. Detection is achieved with an optical fiber probe that is dipped directly into the sample. The probe tip is functionalized with a novel affinity reagent, the MDAC, which recognizes target molecules and generates a fluorescence signal that is then captured by the fiber probe and relayed back to sensitive back-end optics for further analysis.

**Figure 1:**
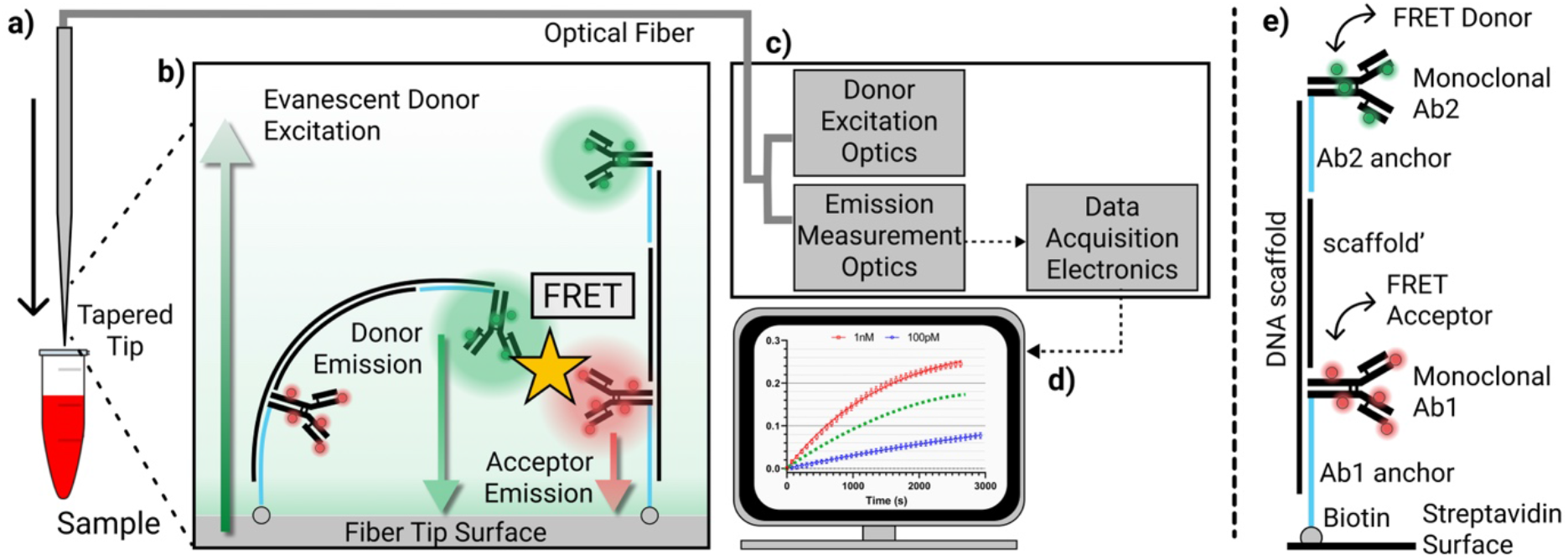
Instant ELISA system overview. Instant ELISA achieves rapid, sensitive detection in complex biological specimens without the need for sample preparation. **a**) A sharply tapered optical fiber probe is dipped directly into the sample. **b**) The surface of the probe tip is functionalized with monolithic dual-antibody clamp (MDAC) molecules, which comprise a pair of fluorescently-labeled antibodies against a common target that are coupled together by double-stranded DNA. Green excitation light travels through the fiber and generates an evanescent field at the tip, exciting the donor-tagged antibody (green) on the MDAC. In the presence of target, many donor-tagged antibodies will be brought into close proximity with acceptor-tagged (red) antibodies that bind to the same target molecule. This gives rise to a Förster resonance energy transfer (FRET) signal, which is coupled back into the fiber and measured in the **c**) optical backend. **d**) The binding rate of the MDAC can then be computed based on the change in FRET ratio. **e**) Structural components of the MDAC molecule.

As with a conventional ELISA, the MDAC achieves specific, sensitive detection by leveraging two antibodies against a common target, and only generating a signal when both are bound simultaneously. In the MDAC design, however, the two fluorophore-labeled monoclonal antibodies are directly linked to each other and to the sensing surface by a DNA scaffold (**Figure 1e**), rather than being added sequentially to the assay with multiple wash steps in between. The two antibodies are each conjugated to 30-nucleotide (nt) DNA oligonucleotides (Ab1-anchor and Ab2-anchor), which are in turn hybridized to a 200-base-pair (bp) DNA scaffold. The DNA scaffold is also hybridized to a fourth oligo (scaffold’) that renders the entire length of the DNA assembly double-stranded. The molecule is tethered to the streptavidin-coated substrate via a biotin modification at the distal end of the Ab1-anchor. The fluorophores on the two antibodies form a donor-acceptor pair for Förster resonance energy transfer (FRET)-based detection, such that donor- and acceptor-tagged antibodies from neighboring MDACs are brought into close proximity upon binding to a common target molecule. This enables rapid detection of binding events without the need for serial incubation and washing steps or the addition of a detection reagent, as is typically the case in conventional ELISA. This design also results in excellent specificity, because the signal is generated only when the antibodies are separated by just a few nanometers, and can be easily generalized to numerous targets by incorporating commercially-available antibody pairs developed for conventional ELISA.

The binding state of the MDAC molecules is interrogated via a previously-reported optical detection hardware setup^27^ that guides excitation light to the MDAC molecules on the probe tip through an optical fiber. The excitation light generates an evanescent field at the sharply-tapered end of the fiber tip (**Fig. 1b**), selectively exciting the surface-bound MDAC molecules. The acceptor fluorophore emission generated by target-bound MDAC molecules is then coupled back into the optical fiber probe and measured by single-photon counting module (SPCM) detectors in the backend. Critically, the exponentially decaying profile of the evanescent field means that auto-fluorescent molecules within the bulk sample volume will not be excited; this results in minimal background fluorescence compared to conventional epifluorescence measurements, enabling continuous interrogation of MDAC-target binding in unprocessed biological samples. By placing the donor green dye on the construct’s distal antibody and the acceptor on the proximal antibody, we minimize dye excitation in the unbound state, leading to reduced photobleaching and increased FRET signal change in the presence of target. Finally, the rate of MDAC-target binding is obtained by computing the FRET ratio, which is defined as the ratio of acceptor emission to the sum of acceptor and donor emissions, and this information is then used to determine the concentration of target in the sample.

### The MDAC construct assembly process

As an initial test, we created an MDAC construct with specificity towards TNFα. We selected two anti-TNFα monoclonal antibodies (clones mAb1 and mAb11 – from here on referred to as clone A and B, respectively) that have been previously shown to bind to be compatible with sandwich ELISA. The MDAC components were then assembled onto streptavidin-coated microbeads in a stepwise fashion (**Figure 2a**). We used N-hydroxysuccinimide (NHS) chemistry to label clone A with the red acceptor dye Alexa Fluor 647 and clone B with the green donor dye Alexa Fluor 546. To assemble the DNA scaffold, we first modified the Fc domain of the dye-labeled antibodies to incorporate an azide group. The Ab1-anchor oligo was synthesized with 3’ biotin and 5’ amine modifications; this 5’ amine group was then labeled with dibenzocyclooctyne (DBCO) using NHS chemistry. We then conjugated the Ab1-anchor with azide-modified clone A via copper-free click chemistry. Ab2-anchor was synthesized and DBCO-labeled in a similar fashion, and then conjugated with azide-modified clone B. We refer to the dye-labeled, oligo-conjugated clone A and B as Ab1 and Ab2, respectively. The scaffold and scaffold’ oligos were then annealed and hybridized to Ab1. We then incubated the streptavidin microbeads with the Ab1-scaffold construct, before finally hybridizing Ab2 with the scaffold.

**Figure 2:**
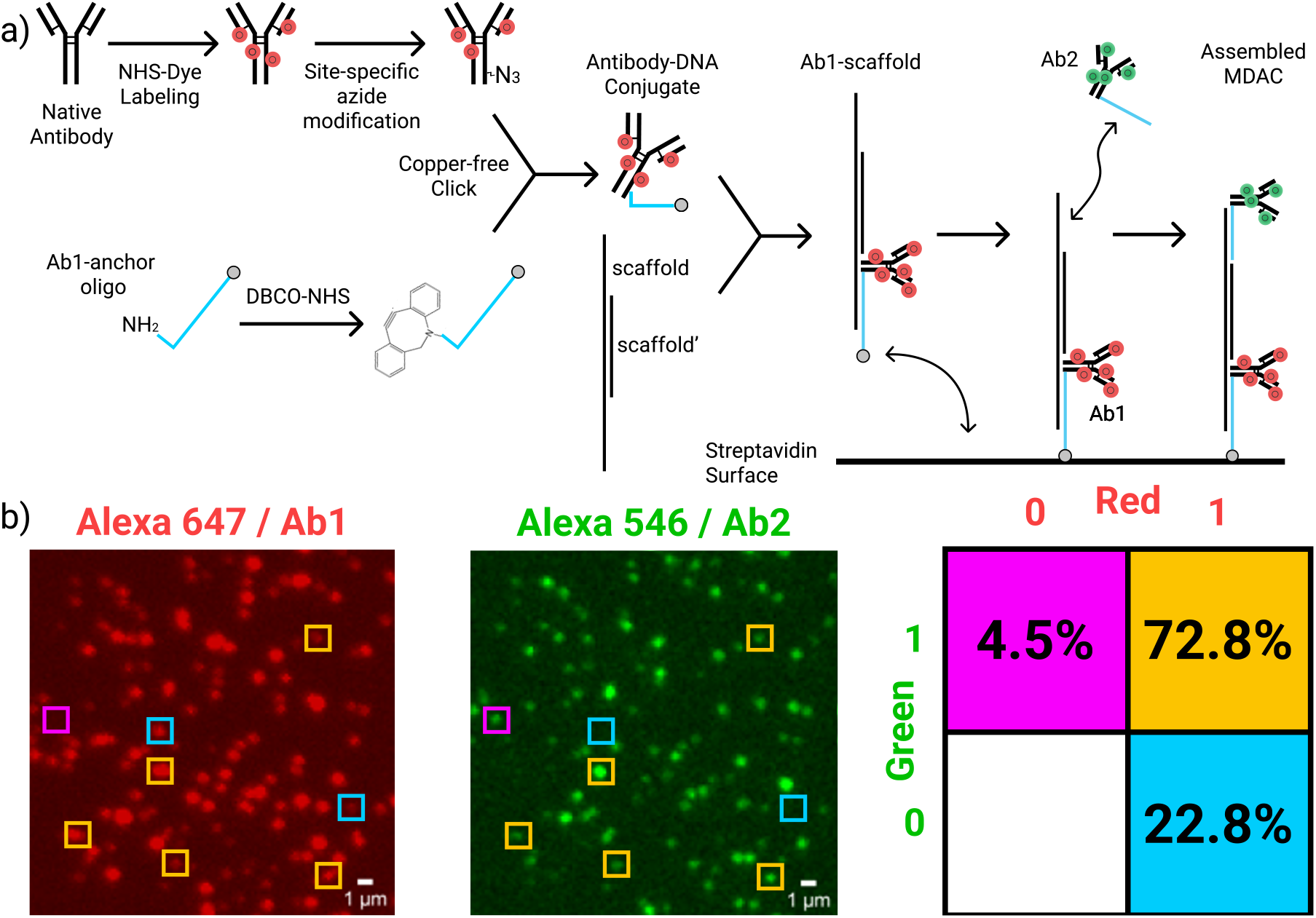
MDAC assembly and characterization. **a**) Process of MDAC assembly. **b**) Single-molecule total internal reflected fluorescence (TIRF) microscopy characterization of the efficiency of MDAC assembly. Constructs were immobilized onto a PEG-passivated coverslip via biotin-streptavidin linkage, and images were recorded in the red (Alexa Fluor 647; 640 nm) and green (Alexa Fluor 546; 532 nm) channels. Colocalization analysis was used to compute the fraction of correctly-assembled constructs, and percentages were computed relative to the fraction of all fluorescent events counted. The boxes in the left and center panels highlight examples of each of the three labeling scenarios shown on the right, and are color-coded to match that righthand panel.

This assembly process was characterized and validated using single-molecule total internal reflection fluorescence (sm-TIRF) microscopy (**Figure 2b**). Using the procedure detailed above, we assembled MDAC constructs onto polyethylene glycol (PEG)-passivated glass coverslips via biotin-streptavidin linkage as previously reported^28^. We determined the fraction of structures that assembled correctly based on spatial colocalization of red and green fluorophores and found that ∼76% of the observed Ab1-scaffold molecules were coupled to an Ab2 molecule, forming a complete MDAC structure. Our data also showed minimal non-specific binding of Ab2 to the surface. Also, by observing the photobleaching signal of individual dyes, we obtained distributions characterizing the number of dye molecules conjugated to per Ab (**Figure S1**). Additionally, we verified via sm-TIRF that the fluorescent dyes do not spontaneously undergo FRET interactions in the absence of target (**Figure S1**). Finally, we validated that we could replicate this same MDAC assembly process on streptavidin microbeads via flow cytometry (**Figure S2**), and the controls performed for that experiment were in agreement with the sm-TIRF data in terms of the efficiency of complete MDAC assembly.

### Assessing MDAC-based detection of TNFα in buffer and complex specimens

We tested the MDAC target response by challenging the MDAC-coated microbeads with a wide range of TNFα concentrations in buffer, incubating them overnight, and analyzing them via flow cytometry (**Figure 3a**). Donor fluorescence sharply decreased and FRET intensity increased over the concentration range 10 pM–1 nM at the donor excitation wavelength (532 nm), while the fluorescence of the acceptor dye changed minimally when directly excited at 638 nm. We computed the FRET ratio (*E*_*pr*_) as *I*_*a*_/(*I*_*a*_ + *I*_*d*_), where *I*_*a*_ is the acceptor emission intensity and *I*_*d*_ is the donor emission intensity, to generate a binding curve for the MDAC (**Figure 3b**). Notably, the signal was reduced at higher target concentrations as individual antibodies started binding to separate target molecules. This conformation does not result in a FRET interaction, and at very high concentrations would ultimately deplete the FRET signal. Interestingly, though the construct was initially designed with the hypothesis that MDAC would bind intramolecularly—*i.e.*, with Abs from the same MDAC structure binding to the same target molecule—rather than intermolecularly—*i.e.*, with each Ab2 cooperating with an Ab1 from a nearby MDAC construct, the sm-FRET data we obtained from MDAC-coated coverslips showed that this is not the case (**Figure S1**). In these sm-TIRF experiments, the streptavidin-coated coverslips were incubated with very low Ab1-scaffold concentrations (50 pM), thereby achieving large intermolecular distances on the order of ∼1 um and rendering target-mediated intermolecular interactions impossible. In this context, no binding signal was observed, indicating that intramolecular binding is not favored, likely due to the rigidity of the double-stranded scaffold. Intermolecular binding does occur on the surface of MDAC beads, which instead are prepared with 100 nM Ab1-scaffold. This form of binding is favored by the flexible linkers joining the Abs to the DNA strands and the flexible PEG linker between the scaffold oligo and the anchoring biotin.

**Figure 3:**
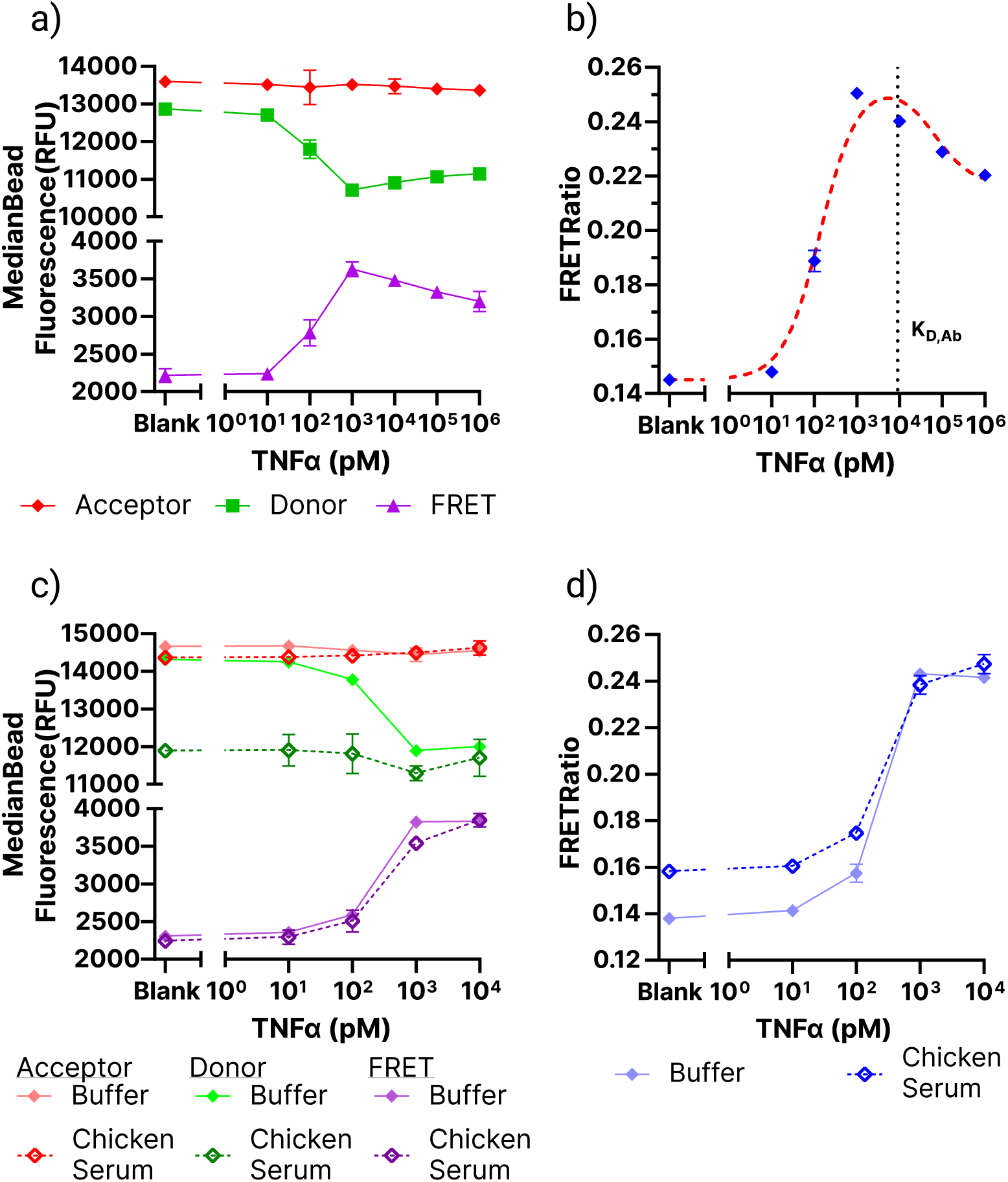
TNFα MDAC characterization. **a**) Flow cytometry-based fluorescence measurements of bead-immobilized MDACs challenged with various TNFα concentrations. **b**) FRET ratios computed from the measurements in panel **a**. The red dashed line shows the predicted binding behavior of this MDAC based on Eq. S1 (**SI Note 1**). **c**) Flow cytometry measurements of bead-immobilized MDACs incubated for 30 minutes with various concentrations of TNFα in buffer and chicken serum. **d**) Calculated FRET ratios from the data in panel **c**. Error bars for all plots represent the standard deviation from three replicates.

We measured an operational dissociation constant, *K*_*D*,0*p*_(*i.e.,* the target concentration at which 50% of the maximum signal is reached) of ∼160 pM. For comparison, the affinities of the constituent antibodies as measured via biolayer interferometry (BLI) were 6.5 nM and 15.1 nM for clone A and clone B, respectively—roughly 70-fold higher (**Figure S3**). This is consistent with our predictions based on a thermodynamic binding model of the construct, which we have described in detail in **SI Note 1**. Briefly, when the target is captured by one antibody clone, it will be able to sweep a restricted volume defined by the length of the DNA scaffold, which limits its entropic state space. Consequently, neighboring antibodies with different clonality will experience a high local concentration of target (*C_eff_*), which enables them to readily bind the target as well. This avidity effect results in slower target dissociation and thus higher effective affinity. Our binding model produced an equation that approximates the binding curve shown in **Figure 3b**. When we fitted this equation to those data, we derived a *C_eff_* of 160 ± 50 nM. We also determined a limit of detection (LOD) of 7 ± 2 pM based on the target concentration at which the signal from the bead-conjugated MDAC is equal to background plus three standard deviations.

We next explored whether this system could achieve this high degree of sensitivity in the complex milieu of chicken serum spiked with human TNFα. To test the construct’s capacity for rapid response, the beads were incubated in serum for only 30 minutes before being isolated, washed, and resuspended in buffer. The procedure was repeated with TNFα in buffer as control. Flow cytometry data from these beads showed a significant increase in FRET emission and concomitant decrease in donor emission between 100 pM and 1 nM TNFα, indicating that the binding kinetics of this MDAC structure are sufficiently rapid to carry out a measurement in only 30 minutes (**Figure 3c**). We also observed a striking decrease in donor fluorescence at lower TNFα concentrations compared to buffer, which resulted in an elevated background signal (**Figure 3d**). We hypothesized that this was attributable to the effects of nucleases present in the serum, which cleave the scaffold and result in loss of the distal donor-labeled antibody. In contrast, the combination of a short DNA linker and steric effects appeared to prevent nucleases from cleaving the proximal, acceptor-labeled antibody from the beads. At higher target concentrations, this nuclease activity is reduced, and we hypothesized that this could be due to steric effects as well— in the presence of target, the structure is brought closer to the bead surface as the two antibodies bind to the target, rendering the DNA backbone less accessible to nucleases. Notwithstanding this degradation effect, we observed roughly equivalent sensitivity after 30 minutes in serum compared to overnight incubation in buffer, with a LOD of 9 ± 11 pM. In an effort to minimize the effects of nuclease degradation, we replaced 20% of the phosphodiester bonds of the DNA scaffold with nuclease-resistant phosphorothioates, and modified the DNA anchor of the distal antibody with interspersed locked nucleic acid (LNA) bases. This modified MDAC exhibited ∼50% reduced degradation in serum (**Figure S4**), and future iterations of MDAC could leverage different synthetic polymers such as L-DNA to fully eliminate the impact of nuclease activity for applications in which such resistance is critical. However, this did not prove necessary for the present work, and all subsequent experiments were performed with entirely natural DNA-based MDACs.

### The MDAC design can be generalized to other protein targets

We next synthesized a new MDAC structure with specificity for a different cytokine, MCP-1, using two monoclonal antibody pairs described in the literature (5D3-F7/10F7 and 5D3-F7/2H5). We repeated the same assembly and verification experiments described above for both pairs (**Figure S5**), but substituted Alexa Fluor 647 with Atto 643 as the acceptor dye due its higher reported quantum efficiency (and thus increased FRET sensitivity) and increased photostability^29^. We ultimately chose to proceed with the 5D3-F7/10F7 antibody pair, which produced a larger FRET signal. We then characterized this MDAC on microbeads by incubating overnight with various concentrations of MCP-1 in buffer (**Figure 4a, b**). We observed comparable performance to the TNFα MDAC, with a concentration-dependent increase in FRET ratio between 10 pM–1 nM MCP-1. This MDAC exhibited a *K*_*D*,0*p*_ of ∼550 pM, which is a considerable improvement over the measured affinity of the individual antibodies, which both had a *K_D_* of ∼6 nM based on BLI (**Figure S5**). We fitted our binding model to the flow cytometry data and calculated a *C*_*eff*_ of 20 ± 5 nM and a LOD of 4 ± 1 pM. We also repeated the rapid 30-minute experiments in chicken serum with the MCP-1 MDAC (**Figure 4c,d**), and observed similar behavior to the TNFα construct. Again, nuclease activity degraded the MDAC scaffold at low target concentrations, but this had negligible impact on the performance of the system, which retained a LOD of 14 ± 6 pM in serum. Thus, the MCP-1 MDAC appears to be highly specific and sensitive, even in the highly complex medium of serum, with sufficiently rapid kinetics and a long enough lifetime to measure picomolar analyte concentrations within 30 minutes.

**Figure 4:**
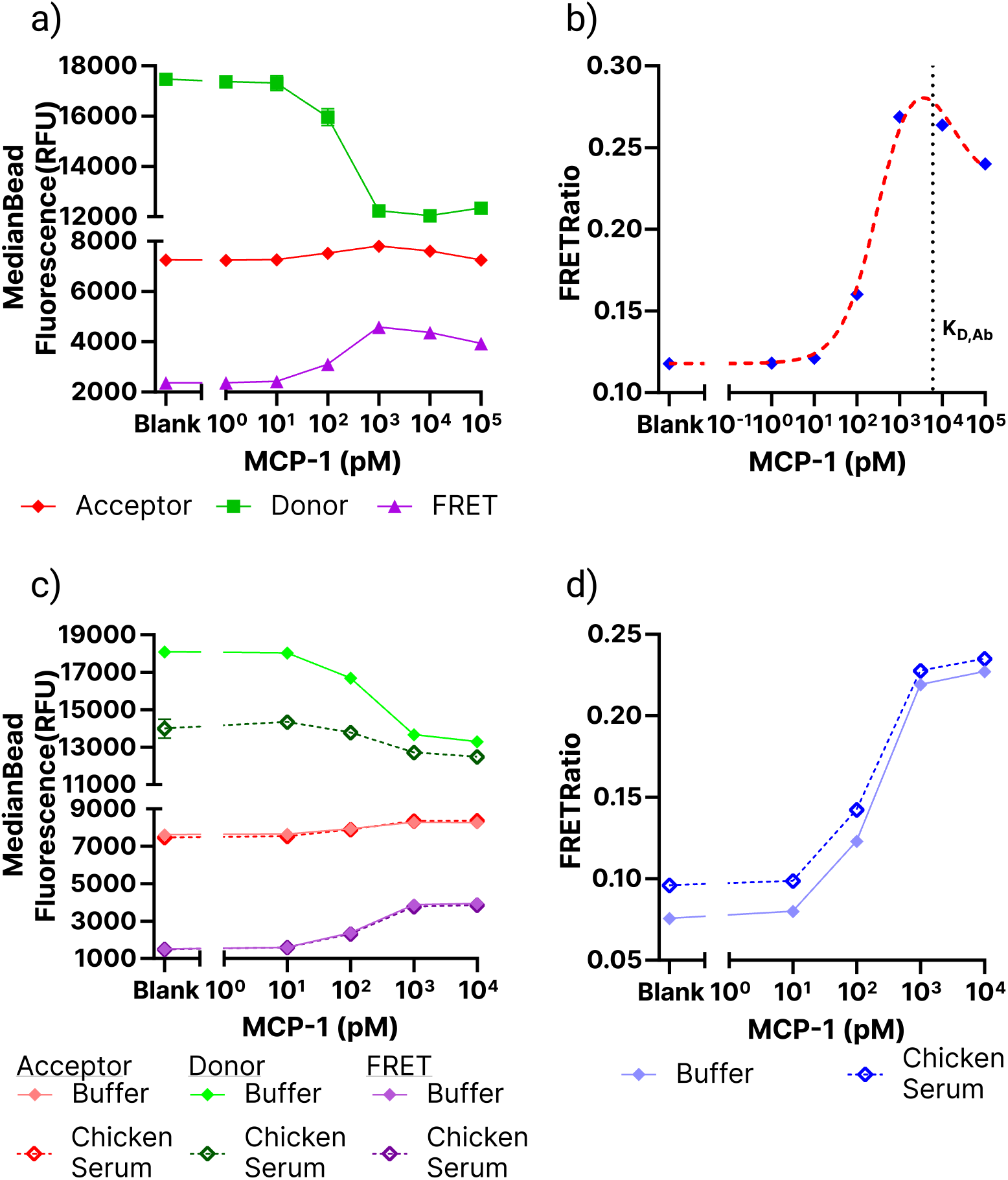
Characterization of the MCP-1 MDAC. **a**) Fluorescence measurements of bead-immobilized MDACs challenged with MCP-1 at various concentrations in buffer and incubated overnight. **b**) FRET ratio at each concentration of MCP-1. The red dashed line shows the predicted binding behavior of this MDAC based on Eq. S1 (**SI Note 1**). **c**) Fluorescence measurements of bead-immobilized MDACs incubated for 30 minutes with MCP-1 in buffer and chicken serum. **d**) Calculated FRET ratios at each MCP-1 concentration in buffer and serum. Error bars for all panels (not visible in some cases) represent the standard deviation of three replicates.

### Instant ELISA rapidly quantifies proteins in complex media with no sample preparation

Having demonstrated the excellent sensing performance of the MDAC affinity reagent, we next incorporated it into our tapered-fiber probe sensor in order to assess its capacity for rapid analyte detection without the need for sample preparation. We began by immobilizing the MCP-1 MDAC construct onto the optical fiber (**Figure S6a**). The surface of the fiber was functionalized via a previously reported three-step protocol^27^, which involves amino-silanization of the oxide surface of the optical fiber probe, followed by conjugation of PEG-biotin via NHS chemistry. The surface was immersed in a solution of neutravidin, which binds to the biotin, and then incubated with the components of the MDAC construct as described above for the streptavidin-coated microbeads. We monitored the fluorescence recorded by the detector to characterize the efficiency of this process and validated the assembly with control experiments (**SI Figure S6b-e**).

We challenged the instant ELISA sensor with a range of concentrations of MCP-1 in buffer (**Figure 5a**). The probe was dipped in the solution, the laser was turned on for 15 s at 75 s intervals, and the donor and acceptor fluorescence intensities reported by the sensor were used to compute the FRET ratio. Sample raw data are shown in **Fig. S7**. We used these data to compute the fractional FRET ratio change, based on the change in FRET ratio compared to the first measurement prior to addition of MCP-1. We repeated this experiment in triplicate for a range of target concentrations. For each replicate, we quantified the rapid initial increase in signal during the first 1,000 s of binding using a linear fit and measured the binding rate in units of fractional FRET ratio change per second. During this initial phase, these linear fits adequately tracked the MDAC binding signal, particularly for low target concentrations (0–250 pM). We saw a significant background FRET ratio change even in the absence of target, but since this increase only occurred while the laser source was turned on, we attributed this drift to asymmetrical photobleaching of the Atto and Alexa dyes. Since Alexa Fluor photobleaches more rapidly than Atto, the remaining direct laser excitation of Atto dyes leads to an increase in FRET signal. Regardless, we found that the observed initial binding rate of the MDAC was proportional to target concentration (**Figure 5a**, **inset**), with a binding rate constant *k* = 9 ± 0.3 × 10^-8^ fractional FRET change s^-1^ pM^-1^. We measured a LOD of 36 ± 23 pM, and the rapid binding kinetics of the MDAC construct enabled us to achieve such sensitive detection within just 1,000 s (∼15 minutes).

**Figure 5:**
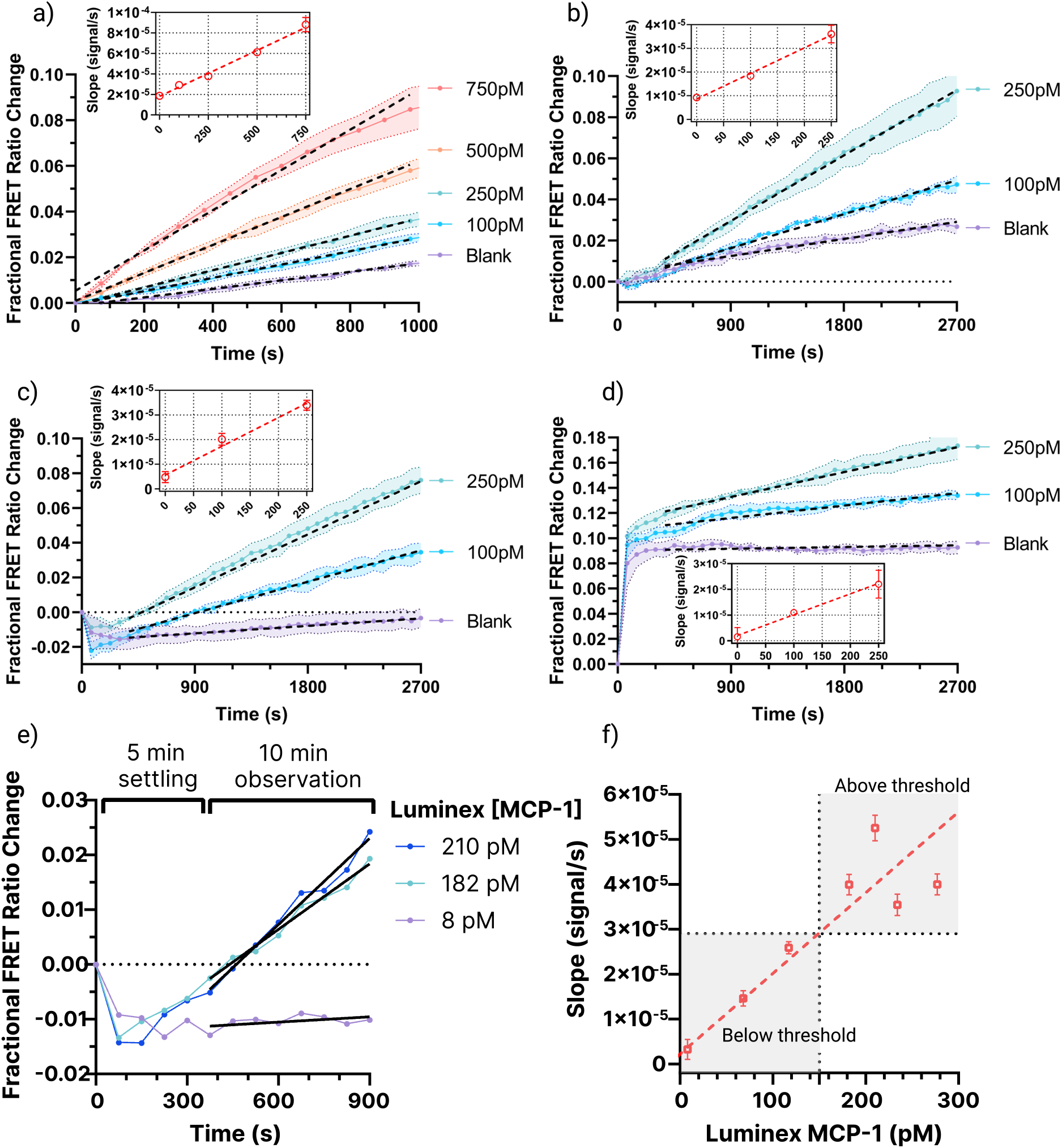
Instant ELISA in buffer and complex media. **a–d**) Characterization of instant ELISA binding signal from MCP-1 in **a**) buffer or chicken **b**) serum, **c**) plasma, or **d**) whole blood. Shaded regions represent standard deviation of at least three replicates. Dashed lines represent linear fits. Insets show best-fit slopes versus target concentration, where error bars represent standard deviation of slopes from at least three replicates. **e**) Instant ELISA data for three of the seven human plasma samples tested (remaining data in **Figure S8**). The probe was allowed to equilibrate for 5 min after immersion in the sample, after which MDAC binding was measured for 10 min. Concentrations in the legend reflect Luminex measurements of each sample. f) The measured MDAC binding rate plotted versus Luminex measurements for each human plasma sample. These data were used to construct a line of best fit correlating MDAC binding rate to MCP-1 concentration, which correlated closely with Luminex measurements (R^2^ = 0.93415). The clinically actionable MCP-1 threshold for heightened CRS risk (150 pM) is shown by the dotted black lines.

Next, we demonstrated rapid quantification of MCP-1 directly in complex media. The sensor was dipped in chicken serum, plasma, and whole blood spiked with small volumes of human MCP-1 to reach concentrations of 100 pM or 250 pM. The laser was turned on for 15 s at 75 s intervals for a total of 45 minutes, during which time the donor and acceptor fluorescence intensities were recorded. The fractional FRET ratio change was computed by comparison to the FRET ratio prior to exposure to complex media (**Figure 5b-d**). Following exposure to the complex media, we noted immediate changes in FRET ratio that were not observed in buffer. For example, in chicken plasma, the FRET ratio consistently decreased immediately after exposure, while the FRET ratio significantly increased after exposure to whole blood. We hypothesized that these sudden changes could be attributed to changes in the chemical environment of the dyes that led to changes in fluorescence intensity. Alternatively, changes in refractive index mismatch between the optical fiber and the solution could alter the profile of the evanescent excitation field and the efficiency of dye emissions coupling back into the optical fiber. Both of these effects could impact the initial FRET ratio, but we found that after this rapid initial change in baseline, the Instant ELISA probe was able to reliably capture the binding signal of MDAC molecules directly in the complex medium. We allowed 5 minutes for the sensor to equilibrate in each medium, and the subsequently-collected data were then fitted to quantify the correlation between binding rate and target concentration (**Figure 5b-d, insets**). The rate constants were obtained through another linear fit, and were very similar to the measured rate constant in buffer (*k*_*serum*_ = 10.8 ± 0.7 × 10^-8^, *k*_*plasma*_ = 11.7 ± 0.7 × 10^-8^, and *k*_*blood*_ = 8 ± 1 × 10^-8^ fractional FRET change s^-1^ pM^-1^). We measured LODs of 8 ± 18 pM for serum, 51 ± 14 pM for plasma, and 126 ± 30 pM for blood.

Finally, we performed an experiment to demonstrate how this assay might be applied in a prognostic setting at the point of care. Cytokine release syndrome (CRS) is a complication that arises in patients receiving CAR T-cell therapy and remains a great challenge for physicians to control.^30,31^ CRS can evolve from manageable symptoms to a severe, life-threatening state in a matter of hours,^32,33^ and rapid prognosis based on cytokine biomarkers could inform timely therapeutic intervention.^34,35^ Studies have shown that MCP-1 holds great promise as a prognostic biomarker of onset of CRS.^35–37^ For example, one study showed that patients who developed CRS grade 4—the most severe form of this condition—exhibited elevated levels of MCP-1 within 36 hours of undergoing CAR T-cell immunotherapy.^38^ The study has specifically proposed an MCP-1 concentration of 150 pM (1.35 ng/mL) as a prognostic threshold for risk of severe CRS.^38^ Currently, these measurements are made with serum or plasma samples via ELISA assays, which have long turnaround times and are of limited value in this time-sensitive context. We therefore tested our instant ELISA system with human plasma samples previously collected for a study of CRS biomarkers in CAR T-cell recipients. This also offered an opportunity to verify that instant ELISA performs as well in human blood samples as in chicken biofluids. Whole blood was collected in EDTA from patients enrolled in a Stanford University IRB-approved study at various timepoints following their CAR T-cell infusion. Within 72 hours of collection, whole blood aliquots were centrifuged at 1,300 × *g* for 10 minutes at 23 °C and the top layer of plasma was pipetted and separated into aliquots of varying volumes. All plasma aliquots were stored in freezer boxes at -80 C until analysis and then thawed before use. MCP-1 concentrations were first quantified using the commercial Luminex immunoassay platform. These measurements were treated as the gold standard for comparison with instant ELISA. The initial FRET proximity ratio was measured prior to exposure to each human sample, after which the probe was dipped in the sample and allowed to equilibrate for 5 minutes. The MDAC binding rate was then observed for 10 minutes (**Figure 5e**). **Figure 5f** shows the binding rate obtained from each human plasma sample plotted against Luminex measurements. Although only a limited number of samples was available for testing, we were able to construct a best-fit line and found good correlation between MDAC binding rate and the Luminex measurements (R^2^ = 0.93415). Although the initial 5-minute settling behavior of the sensor varied from sample to sample, the measured binding rate of the MDAC in human plasma 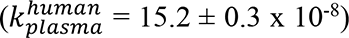 was close to that seen in chicken plasma. Critically, when we set a threshold equal to the clinically actionable concentration of 150 pM, which corresponds to 2.9 × 10^-5^ FRET ratio change s^-1^, we could confidently distinguish between samples falling on either side of this cutoff and did so far faster than time typically required for traditional ELISA tests and commercial immunoassay platforms such as Luminex.

## CONCLUSION

In this work we describe the instant ELISA sensor, which can achieve sensitive quantification of target proteins directly in complex media without the need for sample preparation. We first developed and tested our novel MDAC affinity reagent as a tool for detecting TNFα in buffer and chicken serum, demonstrating that we could achieve a picomolar LOD for this analyte within 30 minutes in complex samples, and proposed an analytical model that approximates the molecule’s binding mechanism. We then showed the generalizability of this design by synthesizing and characterizing a second MDAC for another cytokine, MCP-1. We used this reagent to assemble the complete instant ELISA platform, wherein we coupled the MDAC to an optical fiber sensor that monitors its binding state in real time. We characterized our sensor in buffer and undiluted chicken serum, plasma, and whole blood, and demonstrated that we could consistently achieve sub-nanomolar sensitivity even in complex samples. Finally, we showed that our MCP-1 instant ELISA system could accurately quantify endogenous MCP-1 in undiluted plasma from patients who had undergone CAR T-cell therapy, achieving excellent agreement with Luminex measurements in a fraction of the time (15 min versus several hours).

Several features of the MDAC design contribute to its excellent sensitivity, specificity, and utility for point-of-care detection. The monolithic design allows for all reagents to be tethered to the surface, removing the need for additional reagent addition or wash steps, and the FRET-based sensing mechanism eliminates the need for additional enzyme substrates. And by enforcing a high effective concentration, the DNA linker confers an avidity effect that enables excellent sensitivity—orders of magnitude lower than the *K_D_* of the MDAC’s constituent antibodies. In terms of the sensor hardware, the evanescent field generated by the tapered fiber optic probe enables continuous interrogation of the MDAC in complex biological matrices with minimal background autofluorescence. Although our final MCP-1 measurements were carried out in banked human plasma samples, which involved a minimal amount of sample preparation, we anticipate that similar performance could be obtained with direct measurements in whole blood based on our results from chicken blood. We did not perform such analysis in this work because whole blood samples are rarely banked due to alterations in cytokine levels during storage.^39–45^ Future studies will focus on further clinical validation of this platform with human blood samples tested immediately after collection at the bedside. And since the MDAC utilizes off-the-shelf antibodies as affinity reagents, we believe this platform should be generalizable to any analyte that could otherwise be detected with conventional immunoassays, making our sensor a potentially powerful tool for a range of clinical diagnostic and prognostic assays.

## METHODS

### Reagents

TNFα monoclonal antibodies were purchased from Biolegend (clone mAb1 cat #: 502802; clone mAb11 cat #: 502902). MCP-1 antibodies were purchased from BD Biosciences (clone 5D3-F7 cat #: 551226, clone 10F7 cat #: 555055) and from Biolegend (clone 2H5 cat #: 505902). Recombinant TNFα was purchased from R&D Systems (cat #: 210TA100/CF) and recombinant MCP-1 was purchased from BD Biosciences (cat #: 554620). Oligonucleotides were ordered from Integrated DNA Technologies (all sequences reported in **Table S2**). Oligos were ordered with HPLC purification, except for the scaffold oligo, which was ordered PAGE-purified. Alexa Fluor N-hydroxysuccinimidyl (NHS) esters were ordered from Life Technologies (Alexa Fluor 546, cat #: A20002; Alexa Fluor 647, cat #: A20006) and ATTO 643 NHS ester was purchased from ATTO-TECH GmbH (cat #: AD 643). Site-Click Antibody-Azido modification kits were purchased from Life Technologies (cat #: S20026). Dibenzocyclooctyne-sulfo-N-hydroxysuccinimidyl (DBCO-NHS) ester was purchased from Sigma-Aldrich (cat #: 762040). Anhydrous dimethyl sulfoxide (DMSO) ester was purchased from Sigma-Aldrich (cat #: 276855). Sodium bicarbonate solution 7.5% was purchased from Life Technologies (cat #: 25080094). Amicon Ultra spin dialysis columns were purchased from Millipore Sigma (cat #: UFC503096). Phosphate buffered saline (PBS), 10X (cat #: FLBP399500), sodium chloride, 5M solution (cat #: J60434AE), and Tween 20 (cat #: AAJ20605AP) were purchased from Thermo Fisher Scientific. Bovine serum albumin (BSA) was purchased from New England Biolabs (cat #: B9000S). Normal goat IgG was purchased from R&D Systems (cat #: AB-108-C). Salmon sperm DNA was purchased from Life Technologies (cat #: 15632011). 0.5 M Ethylenediaminetetraacetic acid (EDTA) was purchased from Invitrogen (cat #: 15575020). Dynabeads MyOne Streptavidin T1 were purchased from Thermo Fisher Scientific (cat #: 65601). Biotin was purchased from Invitrogen (cat #: B20656). Mouse Fc-capture biosensor tips were purchased from Sartorius. Chicken plasma (cat #: 502035580) and chicken whole blood (cat #: 502034716) were purchased from Innovative Research. Chicken serum was purchased from Gibco (cat #: 16110082). For reagents related to optical fibers, their preparation and functionalization, see Ref. 27.

### Antibody-oligonucleotide-dye assembly

NHS fluorophores were resuspended in anhydrous dimethyl sulfoxide (DMSO) at 15 mM concentration and stored at -20 °C. Antibodies (0.5 mg/mL) were mixed with sodium bicarbonate solution (0.1 M final concentration). The fluorophores were diluted to 3 mM in deionized water and immediately added to the antibody solutions in different amounts ranging from 1–3% v/v and incubated for 1 h at room temperature. The labeled antibodies were then purified by dialysis (Amicon Ultra 50 kDa MWCO) and the degree of labeling (DOL) was quantified by spectrophotometry using NanoDrop 2000 (Thermo Fisher). The reaction was repeated at larger scale with sufficient dye to obtain a DOL of ∼4-6 fluorophores per antibody. The fluorophore-conjugated antibodies were site-specifically azido-modified using Life Technologies Site-Click Antibody-Azido modification kits following the manufacturer’s instructions. Next, 100 μM amino-modified anchor oligonucleotide was DBCO-functionalized in 0.1 M sodium bicarbonate and 7.5 mM DBCO-NHS. The reaction was incubated overnight at room temperature and then the oligos were purified by ethanol precipitation, vacuum desiccated, and then resuspended at 100 μM in DI water. Finally, the azide-modified antibodies were conjugated with DBCO-functionalized anchor oligos at a 1:10 ratio and incubated overnight at room temperature. The antibody-oligo conjugates were purified from excess oligo by size-exclusion dialysis (Amicon Ultra 50 kDa MWCO).

### Single-molecule total internal reflection fluorescence (sm-TIRF) microscopy

Coverslips were soaked in piranha solution (25% H_2_O_2_ and 75% concentrated H_2_SO_4_) and sonicated for 90 min, followed by five rinses with Milli-Q water. Then, coverslips were soaked in 0.5 M NaOH and sonicated for 30 min, followed by five more Milli-Q water rinses. The slides were then soaked three times in HPLC-grade acetone for 5 min while sonicating, and then dried with N_2_. The dry coverslips were then functionalized with poly(ethylene glycol) silane, MW 5,000 (mPEG-sil). For experiments requiring sample immobilization, a 99:1 ratio w/w of PEG-sil/biotin-PEG-sil was used. The coverslips were incubated in a 25% (w/w) mixture of PEG-sil (or 99:1 PEG-sil/biotin-PEG-sil) in DMSO (anhydrous) at 90 °C for 15 min, then rinsed with water and dried under a N_2_ stream. Imaging chambers (∼8 μL) were constructed by pressing a polycarbonate film with an adhesive gasket onto the coverslip. The surface was then incubated with 12 μL of 0.2 mg/mL (∼200 nM) neutravidin solution for 10 min, and washed twice with 50 μL of 1× PBS (pH 7.5). All experiments were conducted at room temperature (RT; 23 °C).

Next, MDAC reagents were prepared for sm-TIRF experiments. Scaffold and scaffold’ oligos were combined at a final concentration of 20 μM and 30 μM, respectively, in Buffer 1 (1X PBS, 1 M NaCl, 0.1% Tween 20), denatured at 95 °C for 5 min, and allowed to return to RT over 2 h. Ab1 was mixed with excess annealed scaffold (1:10 ratio) at a final concentration of ∼100 nM and 1 μM, respectively, in Buffer 2 (1X PBS, 1 mg/mL BSA, 0.05% Tween 20, 15 μg/mL normal goat IgG, 0.1 mg/mL salmon sperm DNA, 5 mM EDTA, 1M NaCl). The Ab1-scaffold mixture was allowed to hybridize at RT overnight. This solution of Ab1-scaffold was then further diluted in Buffer 2 to a final concentration of 50 pM Ab1. The neutravidin-coated chambers were treated twice with 20 μL of this Ab1-scaffold solution, then washed twice by pipetting 50 μL of Buffer 1 through the chamber. Then, 20 μL of 1 nM Ab2 in Buffer 2 was flowed through the chamber via pipetting and incubated for 60 min to allow hybridization. The coverslip was washed twice by pipetting 50 μL of Buffer 1 through the chamber. Finally, 1 nM target in Buffer 1 was flowed into the chamber via pipetting.

Fluorescence imaging was carried out using an inverted Nikon Eclipse Ti2 microscope equipped with the perfect focus system implementing an objective-type TIRF configuration with a motorized Nikon TIRF illuminator (LAPP) and an oil-immersion objective (CFI Apo TIRF 60× oil immersion objective lens, numerical aperture 1.49). The effective pixel size was 180 nm. 532- and 640-nm lasers were used for excitation (LUN-F XL 532/561/640 Laser Combiner). The laser beam was passed through a multiband cleanup filter (ZET532/640x, Chroma Technology) and coupled into the microscope objective using a multiband beam splitter (ZT532/640rpc-uf2, Chroma Technology). Fluorescence was spectrally filtered with a ZET532/640m-trf (Chroma Technology) filter. For Alexa 546 imaging, fluorescence was further filtered with an ET585/65m (Chroma Technology) emission filter. For ATTO 643 and Alexa 647 imaging, fluorescence was further filtered with an ET705/72m (Chroma Technology) emission filter. Both filters were mounted on a Ti2-P-FWB-E Motorized Barrier Filter Wheel. All movies were recorded onto a 704 × 704 pixel region of a back-illuminated Scientific CMOS camera (Prime 95B, 1.44 MP, Teledyne Photometrics). The camera and microscope were controlled using the NIS-Elements Advanced Research Software Package. For TNFα MDAC photobleaching experiments, excitation was carried out with a power output of 15 mW for the green (532 nm, Alexa 546) laser and 0.9 mW for the red (640 nm, Alexa 647) laser (measured at the objective). Videos were recorded at 7.75 frames per second for four and three minutes for Alexa 546 and ATTO 643, respectively. For FRET experiments, excitation was performed using the 532 nm laser (3 mW). 100 ms snapshots were recorded in the green and red emission channels. For MCP-1 MDAC photobleaching experiments, excitation was carried out with a power output of 12 mW for the green (532 nm, Alexa 546) laser and 12 mW for the red (640 nm, ATTO 643) laser (measured at the objective). For Alexa 546 photobleaching, a three-minute video was recorded at 12.65 frames per second. For ATTO 643 photobleaching, a five-minute video was recorded at 7.72 frames per second. Red-absorbing dyes were always imaged before green-absorbing dyes to avoid unwanted photobleaching. For FRET experiments, excitation was performed using the 532 nm laser (12 mW). 100 ms snapshots were recorded in the green and red emission channels.

Images were analyzed by creating a square ROI in the center of the image (512 × 512 pixels). We observed typically ∼400–800 spots over one field of view (∼92 μm^2^). Fluorescence intensity-time trajectories of individual molecules were extracted from the videos using an algorithm written in MATLAB (MathWorks) defining a diamond shaped region around the centers of the spots captured. A surrounded region was defined to subtract the local intensity background. For the photobleaching analysis, the steps in intensity-time traces were evaluated and counted manually.

### Flow cytometry characterization of MDACs

MyOne Streptavidin T1 beads were washed according to manufacturer’s instructions, and their binding capacity was reduced by incubating 2 mg of beads (200 μL of stock solution) with 2 mL 1 μM biotin solution in Buffer 1 for 30 min on a rotator. Subsequently, the beads were washed three times with Buffer 1 and resuspended in Buffer 1 at 10 mg/mL. Next, Ab1-scaffold hybridization was performed as described above. For experiments with the nuclease-resistant (NR) MDAC, synthesis and assembly was carried out using Ab2-Anchor-NR instead of Ab2-Anchor and Scaffold-NR instead of Scaffold (see **Table S2**). 6 μL of beads were then added to 18 μL of Ab1-scaffold and incubated for 30 min on a rotator, then washed once with Buffer 1 and resuspended in 60 μL of Buffer 2. Ab2 was added to the bead solution to a final concentration of ∼400 nM and incubated for 1 h on a rotator. The MDAC-functionalized beads were then washed with Buffer 1 and resuspended in 60 μL of Buffer 2. For characterization of MDAC binding affinity at equilibrium, 2 μL of MDAC-functionalized beads were dispensed into 200 μL solutions of target protein at a range of concentrations in 1X PBS with 0.1% Tween 20 and incubated overnight on a rotator. Next, the beads were resuspended in 500 uL 1X PBS with 0.1% Tween 20 and immediately imaged on a Sony SH800 FACS machine in analyzer mode. Acceptor and donor fluorescence intensities were obtained by activating the 561 nm and 638 nm lasers and the green and red fluorescence channels. Then, each sample was re-run with the 638 nm laser turned off to obtain the FRET measurements. For the chicken serum and buffer experiments, 2 μL of beads were dispensed into 200 μL of chicken serum or 1X PBS spiked with target protein at different concentrations and 0.1% Tween 20, and incubated for 30 min on a rotator. The beads were then washed twice with Buffer 1, resuspended in 50 μL of Buffer 2, and incubated for 30 min on a rotator. Finally, the beads were resuspended in 1X PBS and analyzed with the Sony SH800.

### Antibody characterization via biolayer interferometry (BLI)

For the characterization of antibody kinetics, a Sartorius Octet Red384 was used with anti-mouse Fc-capture biosensor tips. The tips were rehydrated in DI water for 10 min. The antibodies were diluted to 90 nM in wash buffer (1X PBS, 0.1% Tween 20). The BLI sensor tips were dipped in antibody solution until the BLI shift was around 0.8 nm (∼60 s). The tips were then dipped in wash buffer for 180 s. Different sensor tips were then dipped in dilutions of 80 nM, 60 nM, 40 nM, 30 nM, 20 nM, and 0 nM protein target. The association curve was captured for 300 s, then the tips were moved to blank solution and the dissociation curve was observed for 900 s. ForteBio Octet DataAnalysis software was used to align and fit the data. The off-rate (*k*_*off*_) was determined as the average of the *k*_*off*_ values of each dissociation trace. The fitted association rates (*k*_*obs*_) were then plotted vs target concentration. The on-rate (*k*_*on*_) was determined as the slope of the best fit line, *k*_*obs*_ = *k*_*on*_ [*T*] + *k*_*off*_. Finally, the dissociation constant (*K*_*D*_) was determined as the ratio of *k*_*off*_ to *k*_*on*_.

### Instant ELISA experiments

Optical fibers were tapered and functionalized as previously described.^29^ After preparation, the optical fibers were connected the optoelectronic system^29^ that provides optical excitation and recording of emission intensities. For immobilization of MDACs onto the optical fiber probe, Ab1-scaffold hybridization was performed as described above. Next, the solution was diluted to ∼50 nM Ab1-scaffold in 1X PBS. The neutravidin-treated optical fiber probe was then immersed in the Ab1-scaffold solution and incubated for 1 h. The probe was then dipped in Buffer 1 for 10 minutes to remove unbound Ab1-scaffold. Next, the probe was dipped in 150 nM Ab2 in Buffer 2 for 1 h to allow for hybridization of Ab2 to Ab1-scaffold. The probe was finally washed in 50 μL of Buffer 1 and kept in solution until target quantification experiments were conducted.

For Instant ELISA characterization experiments with buffer, the fiber was dipped in 180 μL of buffer and the laser was turned on for 15s to record the initial signal of the optical fiber probe. Next, 20 μL of target-containing solution was spiked into the tube and the laser was turned on for 15 s at 75 s intervals to record the binding response of the MDAC probes. For characterization in chicken serum, plasma, or whole blood, the medium was spiked with a small volume of MCP-1 to obtain the final desired target concentration. After an initial recording in buffer, the optical fiber was exposed to 400 μL of serum/plasma/blood and the laser was turned on for 15 s every 75 s. For human plasma experiments, after preparation of the fiber probe with MDAC and measurement in buffer, the samples were retrieved from -80 °C storage, thawed, and immediately tested as for the chicken specimens.

The reference Luminex measurements were performed by the Human Immune Monitoring Center at Stanford University - Immunoassay Team. Kits were purchased from EMD Millipore Corporation, Burlington, MA., and run according to the manufacturer’s recommendations with modifications described as follows: H80 kits include 3 panels: Panel 1 is Milliplex HCYTA-60K-PX48. Panel 2 is Milliplex HCP2MAG-62K-PX23. Panel 3 includes the Milliplex HSP1MAG-63K-06 and HADCYMAG-61K-03 (Resistin, Leptin and HGF) to generate a 9 plex. The assay setup followed recommended protocol: Briefly: samples were diluted 3-fold (Panel 1&2) and 10-fold for Panel 3. 25ul of the diluted sample was mixed with antibody-linked magnetic beads in a 96-well plate and incubated overnight at 4°C with shaking. Cold and Room temperature incubation steps were performed on an orbital shaker at 500-600 rpm. Plates were washed twice with wash buffer in a BioTek ELx405 washer (BioTek Instruments, Winooski, VT). Following one-hour incubation at room temperature with biotinylated detection antibody, streptavidin-PE was added for 30 minutes with shaking. Plates were washed as described above and PBS added to wells for reading in the Luminex FlexMap3D Instrument with a lower bound of 50 beads per sample per cytokine. Each sample was measured with single replicates. Custom Assay Chex control beads were purchased and added to all wells (Radix BioSolutions, Georgetown, Texas). Wells with a bead count <50 were flagged, and data with a bead count <20 were excluded. The MCP-1 data collected in Panel 1 was used for comparison with the Instant ELISA measurements.

