## Supplementary Information for "A rapid ELISA platform with no sample preparation requirement"

### SI Note 1: Thermodynamic modeling

The volume confinement provided by the DNA scaffold confers strong avidity for the target. We have proposed an approximate model to describe the expected binding mechanism of MDAC (**Figure S3a**). State U0 describes MDAC in its unbound conformation in the absence of target, where the mean distance of the antibodies generates negligible FRET. When target is present, it will first bind to Ab1 or Ab2 on any given pair of neighboring MDAC molecules (state U1). Equilibrium between U0 and U1 is dependent on target concentration and antibody affinity, which we approximate with a single dissociation constant, $K_D$. At low to moderate target concentrations, the MDAC conformation will readily switch to state S1, where both antibodies are bound to the target. The equilibrium between U1 and S1 is largely independent of target concentration, as the effective concentration of the target with respect to the second binding event is set by the scaffold-linked antibody and its entropic state space. We model this behavior with an avidity factor, $C_{eff}$. State S1 is FRET-competent and generates a binding signal. At higher target concentrations, two different target molecules may bind to each MDAC antibody (state U2). In this state, the dyes do not undergo FRET, and thus the structure does not produce a signal. Thus, we expect a decrease in MDAC signal at high concentrations, resembling that associated with the hook effect in typical immunoassays. The equilibrium between U1 and U2 is determined by target concentration and $K_D$. Due to the existence of two antigen-binding regions on each antibody, the two targets in state U2 may be bound to the antibodies in ways that are unfavorable to further conformational switching. In some cases, however, one of the two targets may be shared between the antibodies. This leads to FRET-competent state S2. The presence of two targets will affect the entropic state space – we model this difference with an adjustment factor, $\alpha$. Boltzmann weights corresponding to the states described above are reported in **Table S1**, and the resulting expression for the total FRET-competent partition, F, is given by:

$$\mathrm{F} = \frac{4\frac{[\mathrm{T}]\mathrm{C_{eff}}}{\mathrm{K_D^2}} + 4\frac{[\mathrm{T}]^2\mathrm{C_{eff}}}{\alpha\mathrm{K_D^3}}}{1 + 4\frac{[\mathrm{T}]}{\mathrm{K_D}} + 4\frac{[\mathrm{T}]^2}{\mathrm{K_D^2}} + 4\frac{[\mathrm{T}]\mathrm{C_{eff}}}{\mathrm{K_D^2}} + 4\frac{[\mathrm{T}]^2\mathrm{C_{eff}}}{\alpha\mathrm{K_D^3}}}. \qquad (1)$$

Boltzmann weight multiplicities are due to the existence of two Fab regions on each antibody, which increases the number of possible binding configurations in each state. In this model, we make several assumptions regarding these configurations. First, we assume that each configuration is thermodynamically equivalent to the others corresponding to the same state. Second, we assume that the FRET competency of each configuration in states S1 and S2 is comparable. Third, we approximate the affinities of both antibodies with one $K_D$ value. Finally, for simplicity, we excluded further binding states such as those involving three or more targets or more than one shared target between the antibodies. These configurations will tend to appear at exceedingly high target concentrations and are expected to be comparatively unfavorable at the concentrations tested in this work.

To explore the implications of this binding model, we began by measuring the affinity of the TNFα antibodies via biolayer interferometry (BLI; **Figure S3b**). The observed on-rates ($k_{obs}$) for different target concentrations were obtained through exponential fits to the BLI sensor association data. The off-rate ($k_{off}$) for each antibody was obtained as the average measured off-rate across all concentrations. The on-rate of each antibody ($k_{on}$) was then obtained as the slope of the best-fit line for Equation 2:

$$k_{obs} = k_{on} \cdot T + k_{off}. \tag{2}$$

The $K_D$ values of the antibodies were then computed via $K_D = \frac{k_{off}}{k_{on}}$, yielding 6.5 nM and 15.1 nM for clone A and B, respectively.

Using the average of these dissociation constants, we then evaluated our model of the MDAC binding curve for a range of $C_{eff}$ values using $\alpha = 10$ (**Figure S3c**). The model shows that for increasing values of $C_{eff}$, $K_{D,Op}$ moves towards lower values (*i.e.*, higher effective affinities). This represents the avidity effect arising from the proximity of the two antibodies, which results in increased sensitivity at lower target concentrations. We also note that for higher values of $C_{eff}$ we observe a larger maximum binding signal, as the binding equilibrium is strongly shifted towards states S1 and S2, disfavoring states U1 and U2. We also evaluated our model with a range of $\alpha$ values where $C_{eff}$ = 120 nM (**Figure S3d**). As expected, this parameter has negligible impact on $K_{D,Op}$, and mostly describes the behavior of the MDAC at high target concentrations.

### SI Note 2: Optical fiber immobilization and controls

We validated the assembly of the MCP-1 MDAC construct onto the optical fiber with two control experiments (**Figure S6a**). In one, Ab1 was omitted to characterize the non-specific binding of the DNA scaffold and Ab2 to the surface; in the other, neutravidin was omitted to verify the specificity of Ab1-scaffold immobilization. We monitored the fluorescence recorded by the detectors in both the green (**Figure S6b**) and FRET channels (**Figure S6c**) at 20-min intervals during steps 4 and 5 of the procedure using three different optical fiber probes. During step 4, after 20 minutes of incubation with Ab1-scaffold, we observed an ~eight-fold increase in fluorescence in the FRET channel compared to background. This signal corresponds to the weak excitation of the red dyes coupled to Ab1 by the green laser. After this initial increase, the fluorescence remained stable for the following 40 minutes. We further characterized this rapid immobilization process by observing the real-time immobilization data for the first two minutes of incubation (**Figure S6d**). We also noted that no fluorescence signal was recorded for the control experiment omitting neutravidin, indicating that immobilization of Ab1 onto the surface of the optical fiber was specifically driven by biotin-neutravidin interaction. During step 5, after 20 minutes of incubation with Ab2, we observed a ~130-fold increase in green fluorescence relative to background. This signal corresponds to the direct excitation and emission of the green dyes coupled to Ab2, after the antibody-oligo conjugate has hybridized onto the Ab1-scaffold constructs immobilized on the optical fiber surface. During this incubation, we observed an increase in the FRET channel as well. As shown by sm-TIRF experiments (**Figure S5e**), this increase is not due to actual FRET interactions between the green and red dyes. Rather, it is due to the leakage of the green dyes in the spectral range of FRET emissions (in flow cytometry experiments, this leakage was subtracted by the instrument automatically). Importantly, due to the nature of the FRET ratio calculation (FRET/(donor + FRET)), this leakage does not impair our ability to perform quantitative measurements. We monitored the hybridization of Ab2 to Ab1 in real-time (**Figure S6e**) and again observed that this process occurred very rapidly. Though we did observe some nonspecific binding of Ab2 to the fiber surface in both control experiments, this was addressed by the final wash step.

**Table S1: Summary of Boltzmann weights for our five-state binding model.**

| State | Boltzmann Weight |
|---|---|
| U0 | $1$ |
| U1 | $4[T]/K_D$ |
| U2 | $4[T]^2/K_D^2$ |
| S1 | $4[T]C_{eff}/K_D^2$ |
| S2 | $4[T]^2C_{eff}/(\alpha K_D^3)$ |

**Table S2: Oligonucleotide sequences used in this work.**

| Name | Sequence |
|---|---|
| Ab1-Anchor | /5AmMC6/GCGAGTTAGGTGGATTAGGGTGATTGAGGC/iSp18/3BioTEG/ |
| Ab2-Anchor | /5AmMC6/TCAACAATAGATAAGTCCTGAACAAGAAAA |
| Scaffold | GCCTCAATCACCCTAATCCACCTAACTCGCCGAATCCTAAACGATTCAGTG GCTAATACAAATCTCATCCACTCACCGCCTAGGCAATCGCTGTCCCAATGC GAAGTCTGAGGTCGGATGCCACGCTCACCGATAGAAGTGTACCTCGGACCA GCGTGACCCTCGCTTCGTTTTCTTGTTCAGGACTTATCTATTGTTGA |
| Scaffold’ | CGAAGCGAGGGTCACGCTGGTCCGAGGTACACTTCTATCGGTGAGCGTGGC ATCCGACCTCAGACTTCGCATTGGGACAGCGATTGCCTAGGCGGTGAGTGG ATGAGATTTGTATTAGCCACTGAATCGTTTAGGATTCG |
| Scaffold-NR | G*CCTCA*ATCAC*CCTAA*TCCAC*CTAAC*TCGCC*GAATC*CTAAA*CGAT T*CAGTG*GCTAA*TACAA*ATCTC*ATCCA*CTCAC*CGCCT*AGGCA*ATCG C*TGTCC*CAATG*CGAAG*TCTGA*GGTCG*GATGC*CACGC*TCACC*GATA G*AAGTG*TACCT*CGGAC*CAGCG*TGACC*CTCGC*TTCGT*TTTCT*TGTT C*AGGAC*TTATC*TATTG*TTGA |
| Ab2-Anchor-NR | /5AmMC6/+TCAA+CAAT+AGAT+AAGT+CCTG+AACA+AGAA+A+A |

/5AmMC6/ indicates a 5’ end amino modification with a C-6 chain linker. /iSp18/ indicates an 18-atom PEG linker. /3BioTEG/ indicates a 3’ end biotin modification with a PEG linker. * indicates a phosphorothioate modification. + indicates an LNA modification.

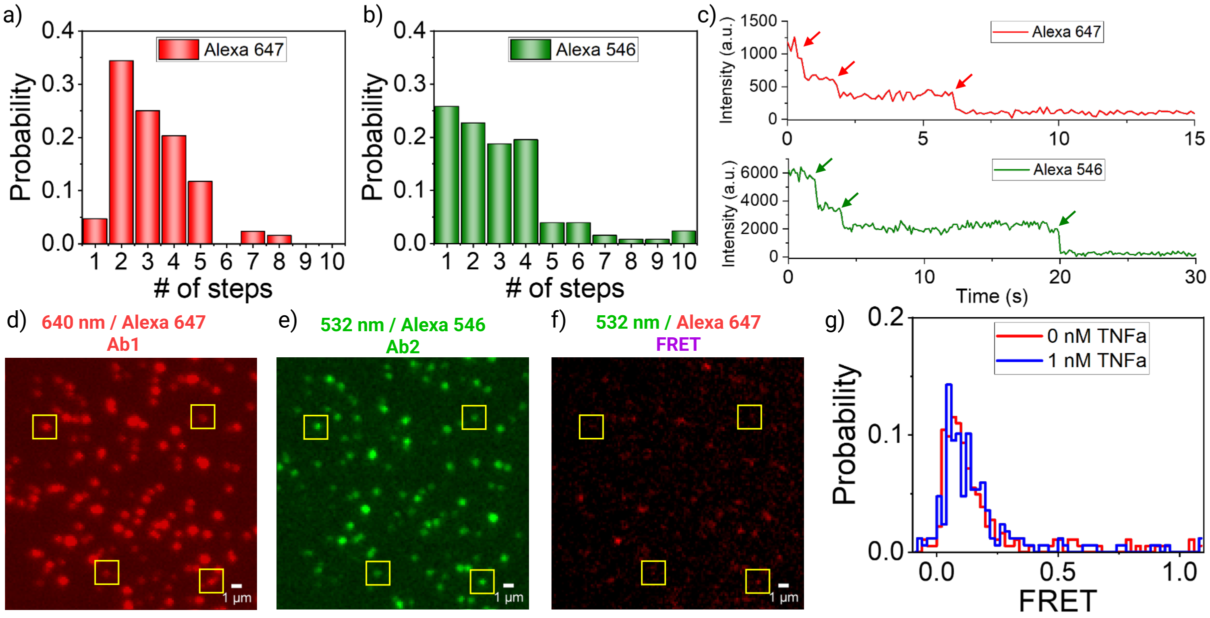


**Figure S1: Additional sm-TIRF assembly data for TNFα MDAC. a–c**) Distribution of the number of **a**) Alexa Fluor 647 dye labels per Ab1 and **b**) Alexa Fluor 546 dyes per Ab2, as measured by **c**) counting the number of intensity steps in a sm-TIRF photobleaching experiment. Arrows highlight representative examples of single-molecule photobleaching events. (**d–g**) MDAC-functionalized surfaces imaged in the **d**) red Ab1 channel, **e**) green Ab2 channel, and f) FRET channel. Yellow squares highlight four correctly assembled structures displaying minimal background FRET interaction. **g**) FRET interactions quantified over a large number of images with and without TNFα, showing that MDAC has minimal background FRET in the absence of target. This background FRET remains unchanged when TNFα is added. This result indicates that intramolecular MDAC binding is not a primary mechanism of FRET signaling, and that FRET signal is instead exclusively generated through intermolecular MDAC interactions. This is because to be able to resolve the fluorescent signal of individual molecules, in these experiments the coverslips were incubated with very low concentrations of Ab1-scaffold to achieve wide spacing (~1 um) between individual MDAC molecules. This large distance renders intermolecular interactions in the presence of TNFα impossible.

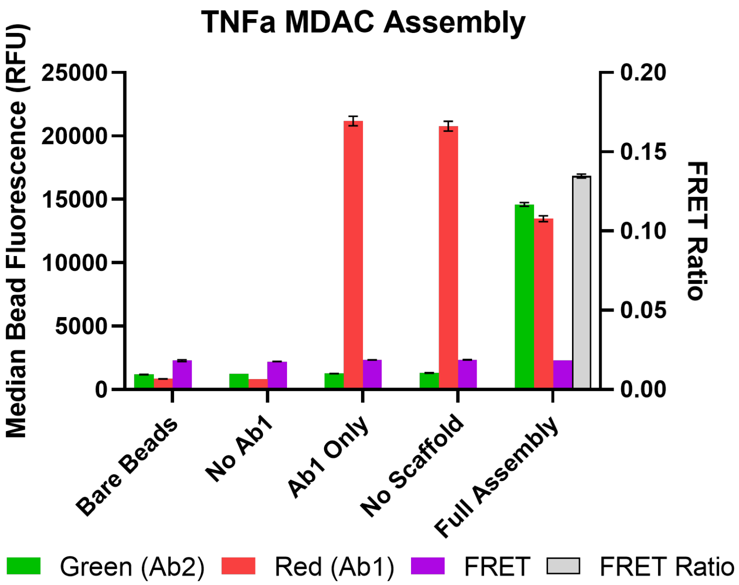


**Figure S2: Flow cytometry-based validation of MDAC assembly.** We analyzed background fluorescence of bare streptavidin beads, beads that omit Ab1 in the assembly procedure, beads with Ab1 only, beads subject to MDAC assembly without the DNA scaffold, and beads with fully-assembled MDACs. Only the final sample shows an increase in both antibody fluorescence channels. The resulting starting FRET ratio is consistent and reproducible.

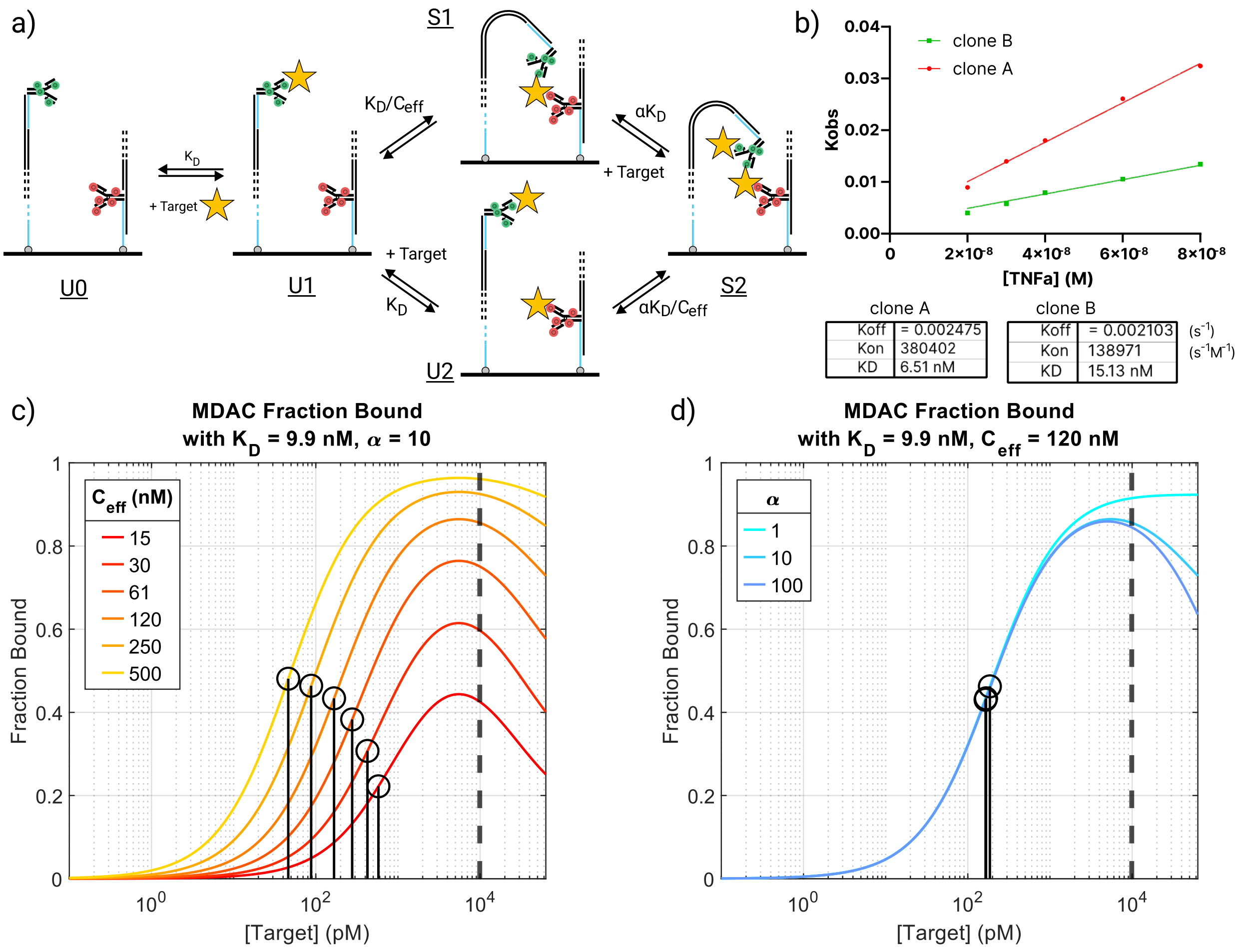


**Figure S3: Thermodynamic model. a**) MDAC five-state binding model and equilibrium relationships. **b**) Determination of $k_{on}$ as the slope of best linear fit to Eq. **Error! Reference source not found.** (error bars are present but not visible) via biolayer interferometry (BLI). Calculation of $K_D$ values for clone A and clone B is shown below. **c, d**) Evaluations of Eq. **Error! Reference source not found.** for different values of **c**) $C_{eff}$ and **d**) $\alpha$. The location of $K_{D,Op}$ for each binding curve is market with ⊙, and the antibody $K_D$ is marked with a dashed line.

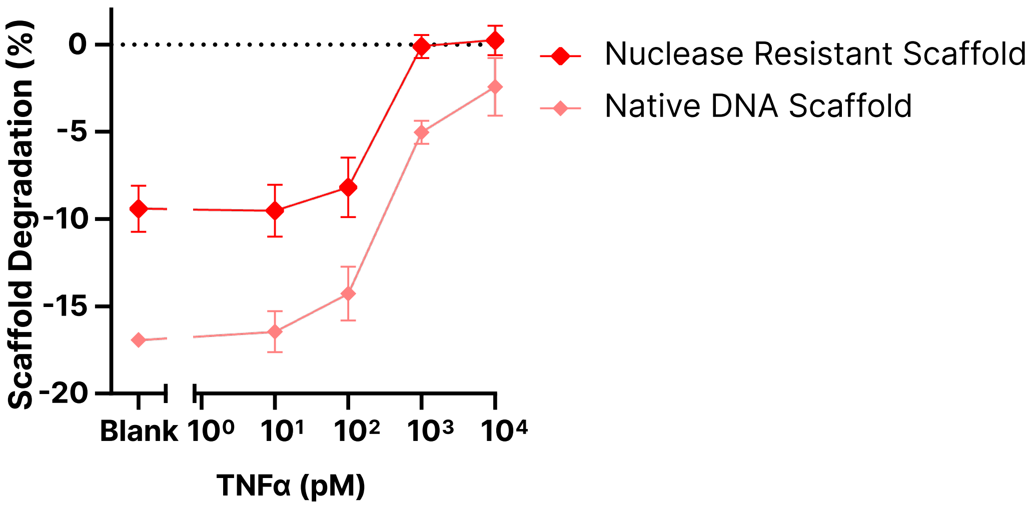


**Figure S4: Nuclease-resistant (NR) DNA scaffold improves MDAC durability in complex media.** MDACs were synthesized using 20% phosphorothioate modifications on the DNA scaffold and 20% locked nucleic acid (LNA) bases in the distal antibody anchor. Degradation of the NR MDAC was compared to the native DNA MDAC after a 30-minute incubation in TNFα-spiked chicken serum. NR MDAC reduced nuclease activity by ~50% at low concentrations, where the scaffold is most susceptible to degradation, and virtually eliminated degradation at higher concentrations.

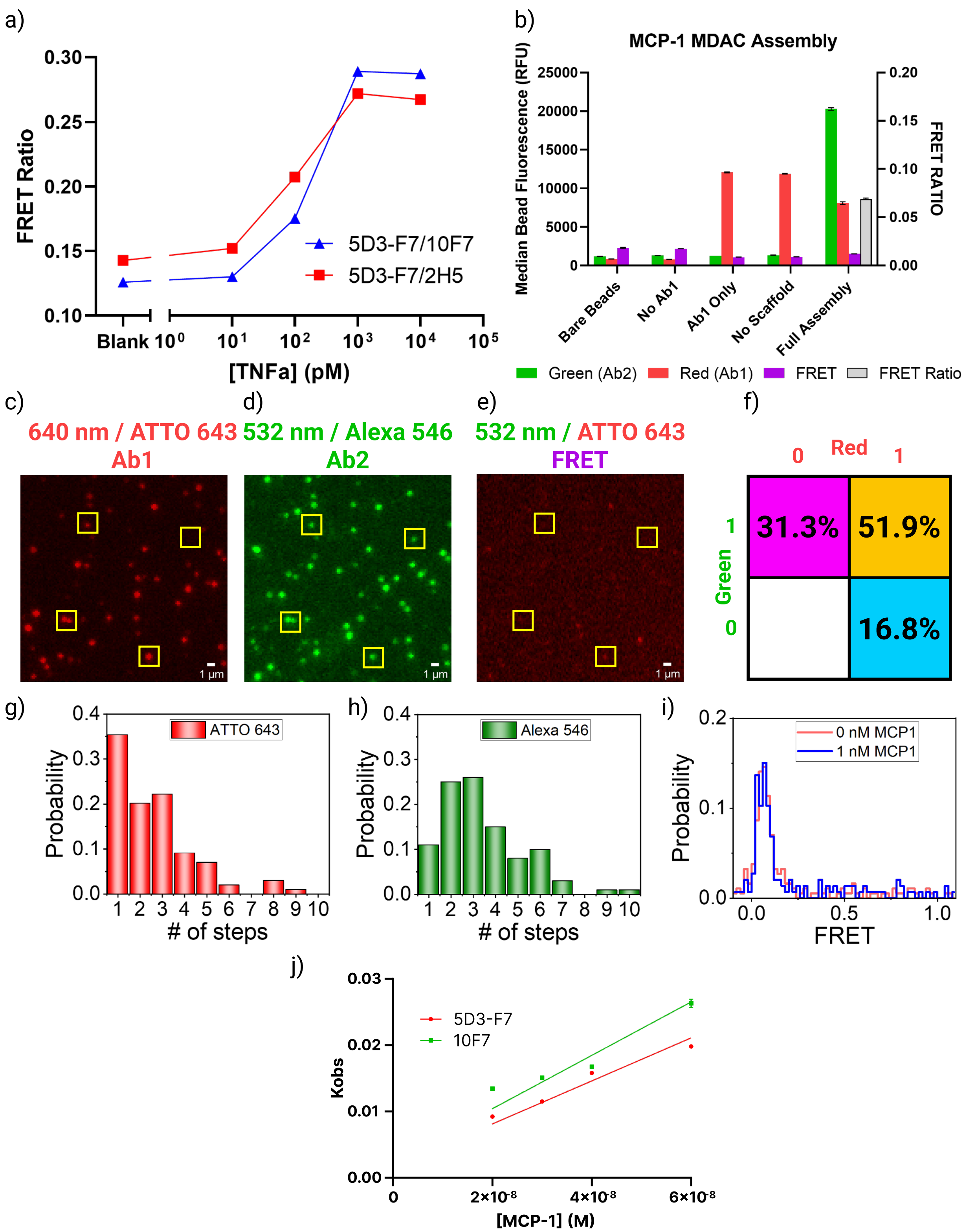


5D3-F7

| | |
|---|---|
| Koff | = 0.001650 |
| Kon | 323431 |
| KD | 5.10 nM |

10F7

| | | |
|---|---|---|
| Koff | = 0.002400 | ($s^{-1}$) |
| Kon | 401600 | ($s^{-1}M^{-1}$) |
| KD | 5.98 nM | |

**Figure S5: Characterization of the MCP-1 MDAC. a**) We evaluated two different pairs of anti-MCP-1 antibodies. The pair 5D3-F7/10F7 was chosen due to the larger binding signal it produced. **b**) MCP-1 MDAC bead assembly was validated via assembly onto streptavidin beads and imaging on a flow cytometer. We analyzed the same set of controls as described in **Figure S2**, but for the MCP-1 antibody pair. The resulting starting FRET ratio was consistent and reproducible. **c-f**) Sm-TIRF characterization of the efficiency of MCP-1 MDAC assembly. Constructs were immobilized onto PEG-passivated coverslips via biotin-streptavidin linkage, and imaged in the **c**) red, **d**) green, and **e**) FRET channels. Examples of correctly-assembled MDACs are highlighted by yellow boxes. **f**) Colocalization analysis was used to compute the fraction of correctly-assembled constructs relative to all fluorescent events counted. **g**) Distribution of the number of Atto 643 dyes per 5D3-F7 antibody and **h**) Alexa Fluor 546 dyes per 10F7 antibody as measured by a sm-TIRF photobleaching experiment. **i**) FRET interactions quantified over a large number of images with and without MCP-1 show minimal background FRET in the absence of target. This background FRET remains unchanged when target is added, again confirming the lack of intramolecular binding occurring and the need for closer spacing between MDACs to produce an intermolecular FRET signal. **j**) Determination of $k_{on}$ via BLI, based on the slope of best linear fit to Eq. **Error! Reference source not found.**. Error bars are present but not visible. Bottom shows calculation of $K_D$ values for 5D3-F7 and 10F7.

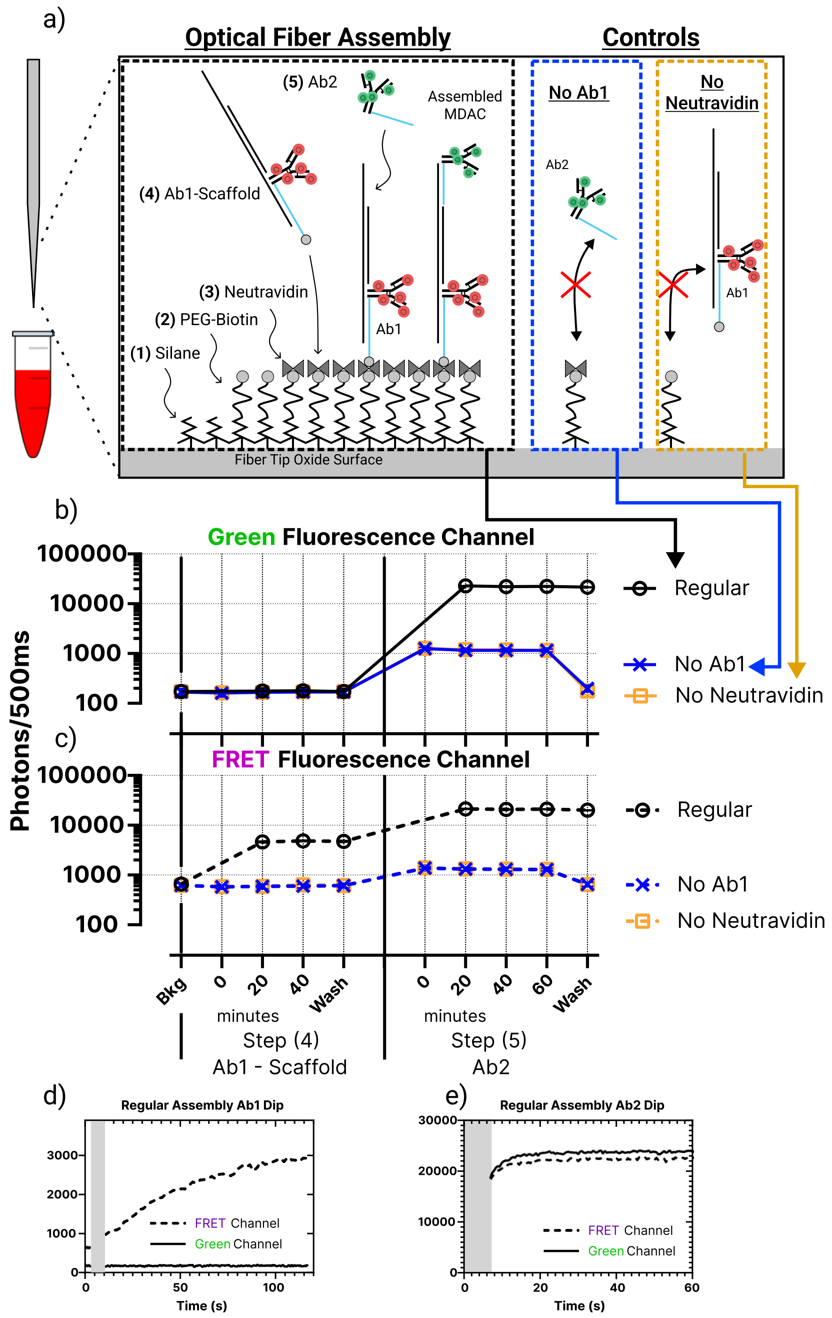


**Figure S6: Immobilization of the MCP-1 MDAC onto optical fiber**. a) Scheme for immobilization of MDAC onto the optical fiber probe. This process entails: **(1)** amino-silanization of the optical fiber tip; **(2)** biotinylation/passivation of the surface with PEG-biotin; **(3)** incubation of the surface with neutravidin; **(4)** incubation with the Ab1-scaffold construct; **(5)** hybridization with Ab2. Control experiments omitted Ab1 or neutravidin. (**b, c**) Fluorescence from the **b**) green and **c**) red channels measured via instant ELISA during experimental verification of steps (**4**) and (**5**). Fluorescence intensity was briefly monitored at 20-minute intervals during the assembly process. Error bars (not visible) represent the standard deviation of three experiments. (**d, e**) We monitored **d**) the first two minutes of step (**4**) and **e**) the first minute of step (**5**) incubation in real time for one fiber. Grayed-out portions of these graphs indicate the time required to dip the fiber in the solution and initiate the measurement.

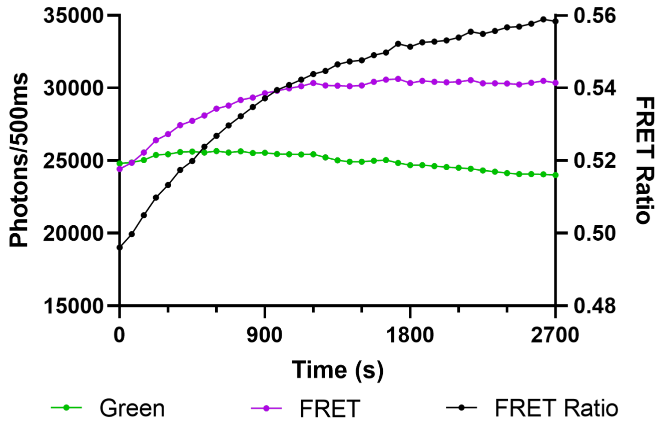


**Figure S7: Raw detector output and computed FRET ratio**. Example of unprocessed data from instant ELISA for detection of 750 pM MCP-1 spiked into buffer. Plot shows the mean photon count per 500 ms measured in each 15 s interval for a 45-minute experiment, as well as the FRET ratio computed based on these measurements. Although the laser power drifted throughout the duration of the experiment, the ratiometric FRET readout was able to capture the binding signal of the MDAC probes in real time.

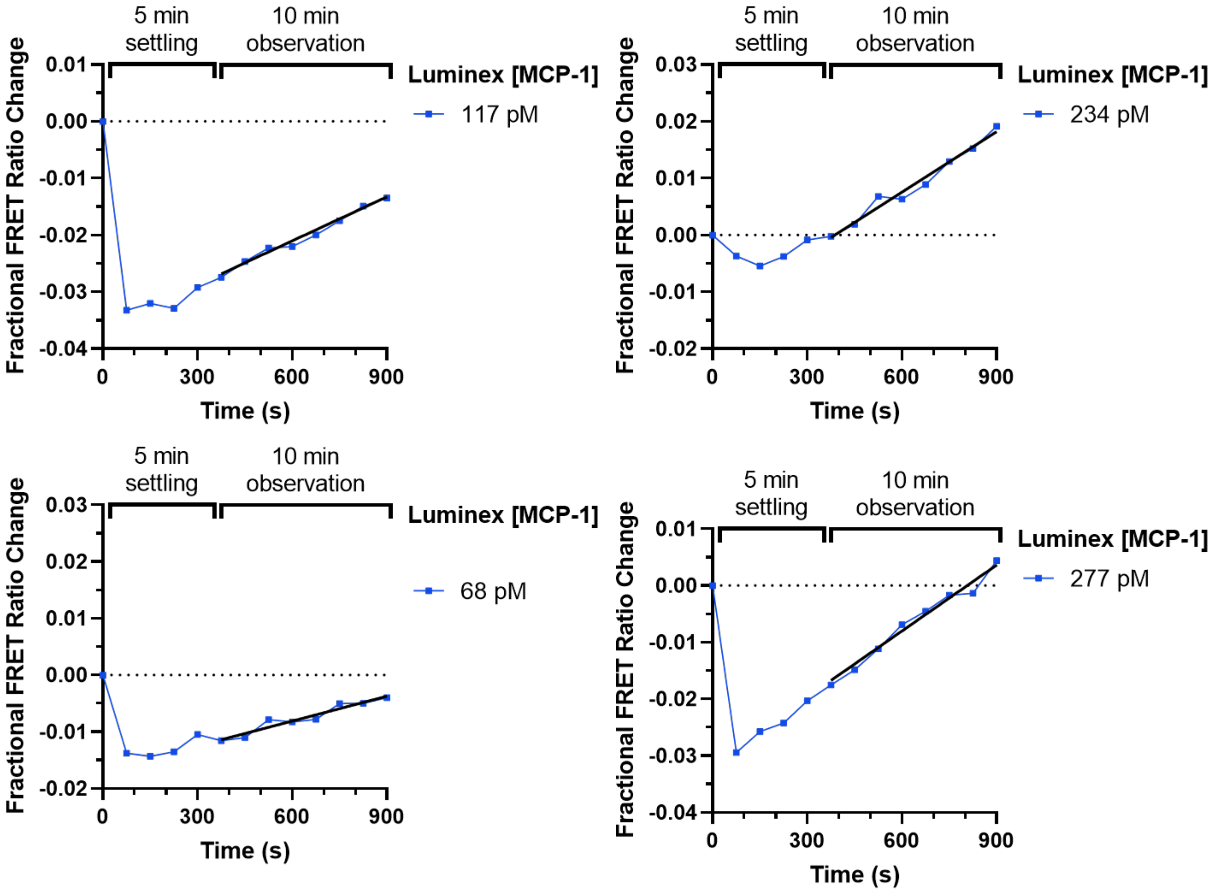


**Figure S8: Performance of instant ELISA for MCP-1 in human plasma.** Data are for four out of the seven human plasma samples tested in this work. The probe was allowed to equilibrate for 5 min after immersion in the sample, after which the MDAC binding rate was measured for 10 minutes. Reference Luminex measurements of MCP-1 concentration for each sample are shown in the legends of each plot.
